# Global mapping of protein localization in vegetative *Dictyostelium discoideum* by subcellular spatial proteomics

**DOI:** 10.64898/2026.09.28.755154

**Authors:** Statton Tinker, Yu-Ping Poh, Katarzyna Dabrowska, Ritin Sharma, Patrick Pirrotte, Jeremy G. Wideman

**Affiliations:** Center for Mechanisms of Evolution, Biodesign Institute, School of Life Sciences, Arizona State University, USA; Integrated Mass Spectrometry Shared Resource, City of Hope Comprehensive Cancer Center, Duarte, CA USA; Division of Early Detection and Prevention, Translational Genomics Research Institute, Phoenix, AZ USA

**Keywords:** Subcellular spatial proteomics (SSP), LOPIT-DC, *Dictyostelium discoideum*, protein localization

## Abstract

*Dictyostelium discoideum* is a genetically tractable amoebozoan model organism widely used to study conserved eukaryotic cellular processes, yet a comprehensive map of its subcellular proteome has been lacking. Here, we combined subcellular fractionation with label-free mass spectrometry to resolve a global protein localization map for *D. discoideum*. We detected 6,337 proteins (~50% of the predicted proteome) and generated protein abundance profiles across 10 subcellular fractions. Using a curated marker set and a support vector machine classifier with a median cutoff, we assigned high-confidence localizations to 3,169 proteins across 17 subcellular compartments. Independent validation using sequence-based targeting predictions strongly supported the compartment assignments. Together, this study provides the first global subcellular localization map of the *Dictyostelium* proteome, establishing a foundational resource for functional, cell biological, and evolutionary analyses. We are currently preparing manuscripts that further analyze (i) the membrane trafficking system of *Dictyostelium*, including detailed investigation of the contractile vacuole, Jotnarlogs, and patchy proteins, and (ii) the composition of the *Dictyostelium* peroxisome, with a focus on the subcellular localization of sterol biosynthesis enzymes. Investigators interested in using these data are encouraged to contact us prior to publication to avoid overlap and to facilitate coordinated and collaborative use of this resource.

## Introduction

The amoebozoan *Dictyostelium discoideum* (*Dictyostelium* henceforth) is a free-living soil amoeba, typically ranging from 10 to 20 µm in diameter, with a haploid genome consisting of six chromosomes that encode approximately 12,500 proteins (Eichinger et al., 2005). First described in 1935, *Dictyostelium* gradually gained prominence as a model organism in cell biology as an alternative to more traditional systems such as yeast (Raper, 1935; Torija et al., 2006). Because amoebozoans branch sister to Obazoa (a large multi-kingdom assemblage that includes fungi, animals and their closest protist relatives), *Dictyostelium* shares many orthologous proteins and processes with animal cells (Baldauf et al., 2000). For example, *Dictyostelium*’s amoeboid morphology makes it particularly valuable for exploring processes such as chemotaxis, phagocytosis, endocytosis, and host-pathogen interactions (Jauslin et al., 2021; Nichols et al., 2015; Vines and King, 2019). In recognition of its biomedical relevance, the National Institutes of Health (NIH) designated *Dictyostelium* as one of eight non-mammalian model organisms for studying human cell biology and disease mechanisms (Martín-González et al., 2021). Research on *Dictyostelium* has provided insights into immune system function, neurodegenerative disorders, and cancer, particularly in relation to cell motility and migration (Frej et al., 2017; McLaren et al., 2019; Myre, 2012). Its tractable genetics and ease of manipulation have enabled researchers to study complex cellular processes that are difficult to study in other systems (McLaren et al., 2019; Myre, 2012). Genomic and proteomic data are curated and made publicly accessible through *dictyBase*, an essential resource for the *Dictyostelium* research community (Chisholm, 2006).

Beyond its utility as a model organism, *Dictyostelium* has a complex life cycle that includes a multicellular phase (Annesley and Fisher, 2009). In response to starvation, individual *Dictyostelium* cells release the chemoattractant cyclic adenosine monophosphate (cAMP), which attracts thousands of neighboring cells and triggers aggregation into a multicellular structure within hours (Bonner, 1944; Konijn et al., 1967). This rapid and highly coordinated response progresses through distinct developmental stages, including aggregation, mound formation, and slug migration, ultimately culminating in the formation of a fruiting body elevated above the substrate by a stalk composed of differentiated, apoptotic cells (Chisholm and Firtel, 2004).

Despite its widespread use, a cell-wide subcellular proteomic analysis has not yet been performed using *Dictyostelium*. To identify protein localization across the cell, we applied a subcellular spatial proteomics workflow to generate a global protein localization map. This approach combines cell fractionation with mass spectrometry to resolve proteins into subcellular clusters and has been successfully applied to diverse eukaryotes (Barylyuk et al., 2020; Chen et al., 2025; Chisholm et al., 2026; Christoforou et al., 2016; Dunkley et al., 2004; Foster et al., 2006; Geladaki et al., 2019; Guérin et al., 2023; Hammond et al., 2025, 2026; Imhak et al., 2017; Jirsová et al., 2025; Moloney et al., 2023; Nightingale et al., 2019; Thul et al., 2017; Zítek et al., 2022). Using this strategy, we detected and predicted the steady-state localization of 3,169 proteins across 17 subcellular compartments, providing a reference dataset for protein localization in vegetative *Dictyostelium*.

## Results/Discussion

### Subcellular Fractionation and Proteomic Profiling of Dictyostelium discoideum

To capture the subcellular organization of *Dictyostelium* proteins under vegetative conditions, axenically grown cells were fractionated by differential ultracentrifugation and analyzed by mass spectrometry. Approximately 10^7^ axenically grown cells were gently lysed using three complementary methods designed to disrupt the plasma membrane while preserving organelle integrity (Figure 1A). The resulting lysates were separated into 10 distinct subcellular fractions by differential ultracentrifugation. Successful enrichment of major organelles was confirmed by Western blotting using antibodies against established marker proteins for the contractile vacuole (vacuolar ATPase subunit A)(Liu and Clarke, 1996), mitochondria (porin A)(Troll et al., 1992), nucleus (H3 histone), and endoplasmic reticulum (PDI) (Gilbert, 1997) (Figure S1). Clear separation of these markers across fractions allowed us to move forward with mass spectrometry.

**Figure 1.**
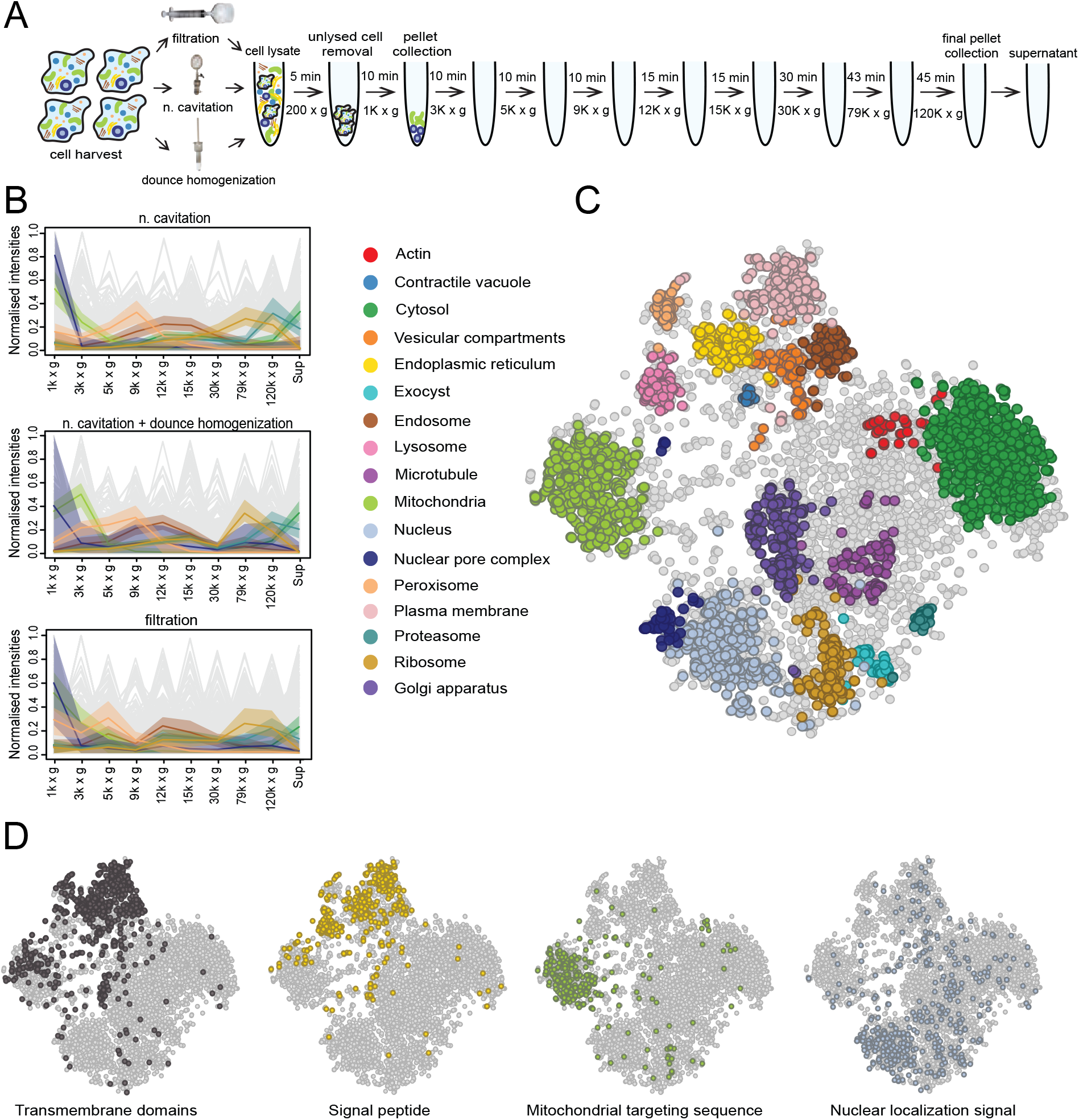
Subcellular spatial proteomics localizes 3,169 proteins to 17 compartments in vegetative *Dictyostelium* cells. **A**. Gentle lysis followed by differential centrifugation generated 30 fractions across three independent lysis experiments. **B**. Normalized abundance profiles were generated through mass spectrometry analysis. The highlighted clusters include the Golgi apparatus, ribosome, endosome, nuclear pore complex, peroxisome, cytosol, and mitochondria, and were selected for clarity to illustrate distinct fractionation profiles across fractions. Grey lines represent all remaining protein abundance profiles. **C**. A supervised support vector machine (SVM) clustering algorithm, trained using 252 marker proteins selected from the curated marker set (Table S1), was used to predict protein localizations, represented on a t-SNE plot. Predictions shown are based on application of a median SVM score cutoff to retain high-confidence assignments. **D**. Predicted targeting features were mapped onto the subcellular atlas. Signal peptides identified by SignalP 6.0 (Teufel et al., 2022), transmembrane proteins predicted by DeepTMHMM v1.0.24 (Hallgren et al., 2022), mitochondrial-targeted proteins predicted by MitoFates v1.2 (score > 0.9) (Fukasawa et al., 2015), and nuclear localization signals predicted by NLStradamus r.9, (score > 0.95) (Nguyen Ba et al., 2009).

Label-free mass spectrometry analysis of the 10 fractions identified 6,337 proteins, corresponding to ~50% of the ~12,500 predicted proteins in the *Dictyostelium* predicted proteome (Eichinger et al., 2005). This represents approximately 89% of genes expressed during vegetative growth using a ≥30-read transcriptomic threshold (Rosengarten et al., 2015). Proteins not detected are attributable to a combination of stage-specific protein expression (Banu et al., 2026) and the inherent detection limits of mass spectrometry-based proteomics (Jiang et al., 2024). As a result, proteins expressed at low abundance or primarily outside the vegetative stage may not have been identified. Protein abundances were normalized, missing values were imputed, and protein abundance profiles across the 10 fractions were generated for each protein detected by mass spectrometry (Figure 1B). These profiles were subsequently visualized by t-distributed stochastic neighbor embedding (t-SNE), revealing clear clustering of proteins (Figure 1C).

### Subcellular spatial proteomics and support vector machine classification localizes >3,000 proteins to 17 compartments in *Dictyostelium*

To define subcellular compartments, we compiled a high-confidence training set of marker proteins with either direct experimental evidence in *Dictyostelium* or strong inferred homology to validated markers in other eukaryotes (Table S1). A subset of 252 of these markers was used to train a support vector machine (SVM) classifier on the fractionation profiles. Application of the trained SVM with a median probability cutoff assigned subcellular localizations to 3,169 proteins (Figure 1C), thereby providing the first predicted localization for thousands of previously unannotated *Dictyostelium* proteins. The clustering structure is consistent across imputation schemes (Figure S2A–C), indicating that missing value handling has minimal impact on the overall organization of the subcellular localization profiles. Using an unsupervised k-means approach with k = 17, the algorithm partitioned the data into 17 clusters (Figure S2D) that broadly match the major subcellular structures in the dataset. Overall, the SVM-based assignments show strong agreement with the k-means clusters, indicating consistent structure between the two methods. The reliability of the SVM assignments was assessed using sequence-based prediction tools (Figure 1D). Mitochondrial targeting sequences (predicted by MitoFates v1.2) were overwhelmingly enriched in the mitochondrial cluster (Fukasawa et al., 2015). Signal peptides predicted by SignalP 6.0 were predominantly associated with compartments of the membrane trafficking pathway, including the endoplasmic reticulum, Golgi apparatus, lysosomes, endosomes, and plasma membrane (Teufel et al., 2022). Transmembrane domains predicted by DeepTMHMM v1.0.24 were enriched in all membrane-bound organelle clusters but absent from the nuclear and cytosolic clusters, as expected (Hallgren et al., 2022). Nuclear localization signals (NLStradamus r.9) were strongly enriched within the nuclear cluster, providing independent support for the assignment of nuclear proteins (Nguyen Ba et al., 2009). The strong connection between fractionation-based SVM predictions and independent sequence-feature analyses supports the accuracy of the subcellular map. This comprehensive resource expands the currently annotated proteome of *Dictyostelium* and provides a reliable foundation for future functional and evolutionary studies.

### Global fractionation refines the *Dictyostelium* mitochondrial proteome

Comparison of the mitochondrial cluster generated in this study with the mitochondrial compendium reported by (Freitas et al., 2022) revealed substantial agreement while highlighting important differences between subcellular spatial proteomics and traditional organelle-enrichment approaches. Of the 936 proteins included in the Freitas et al. mitochondrial compendium, 809 were detected in our dataset. Unlike Freitas et al., who purified mitochondria by density-gradient centrifugation prior to mass spectrometry, our workflow fractionated the entire cellular proteome using complementary lysis methods and differential centrifugation. Consequently, every detected protein was evaluated relative to all other cellular compartments, allowing proteins to be assigned according to their complete fractionation profiles rather than their enrichment within an isolated mitochondrial fraction.

Removing the SVM score cutoff increased agreement with the Freitas et al. compendium while also revealing additional mitochondrial candidates unique to our dataset. At the median SVM score cutoff, 531 proteins were shared between our mitochondrial predictions and the Freitas et al. compendium, while 17 proteins were uniquely predicted by our SVM classifier (Figure 2A). Removing the median score cutoff increased the overlap to 672 proteins and identified 92 additional SVM-predicted mitochondrial proteins absent from the Freitas et al. compendium (Figure 2B). Among the 809 proteins from the Freitas et al. compendium detected in our dataset, 672 were predicted as mitochondrial, 109 localized to other predicted compartments including the peroxisomal protein MfeA (Matsuoka et al., 2003), and 28 localized adjacent to the mitochondrial cluster. The 28 adjacent proteins were retained after inspection of their fractionation profiles, resulting in 700 proteins supported by both datasets. These observations indicate that proteins associated with other organelles can co-purify during mitochondrial enrichment but can be distinguished by global subcellular fractionation. The 92 SVM-predicted proteins absent from the Freitas et al. compendium were subsequently evaluated based on their positions relative to the mitochondrial cluster, resulting in 66 additional proteins being retained in the final curated compendium.

**Figure 2:**
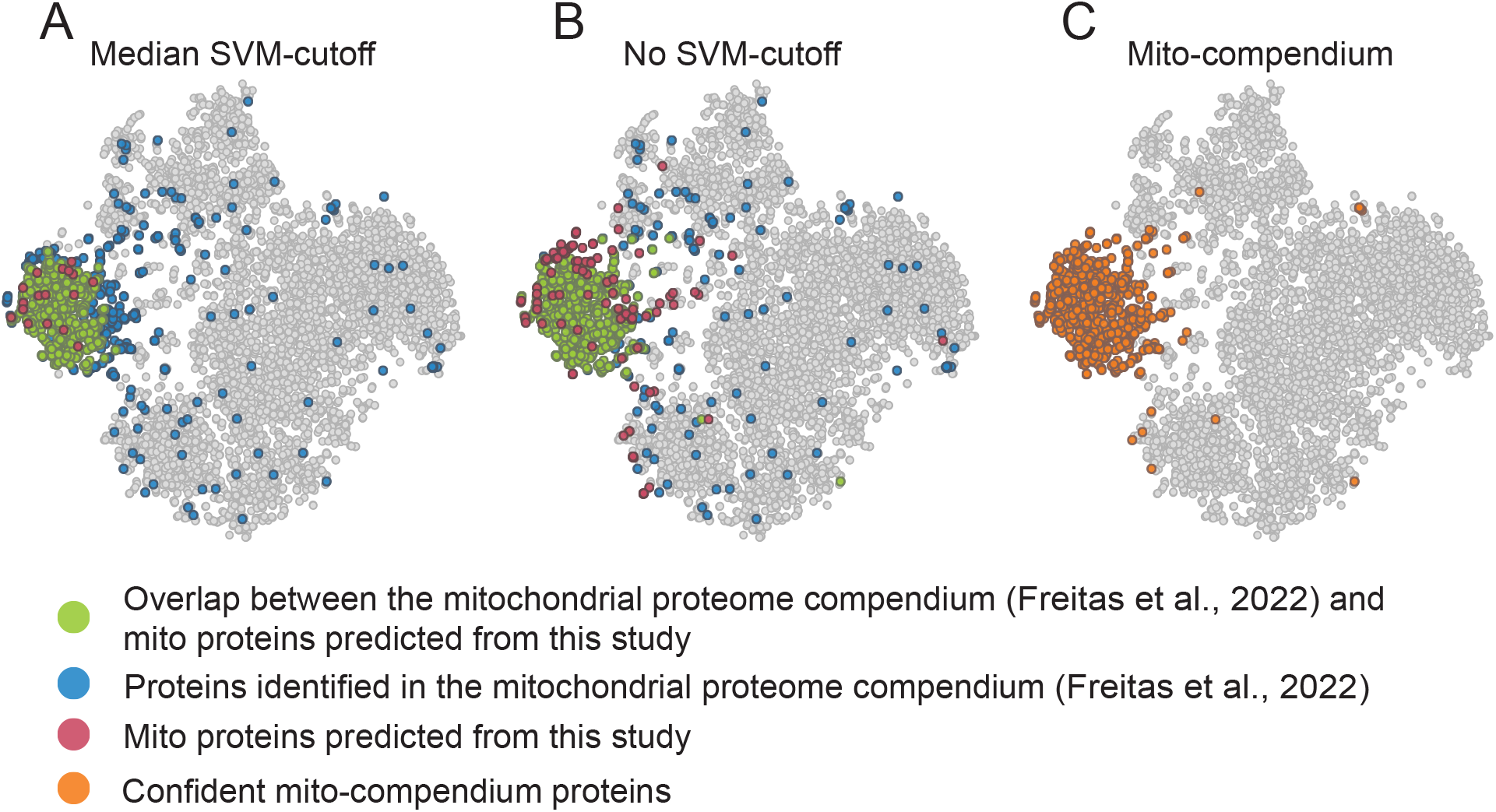
Comparison of SVM-predicted mitochondrial proteins with a published mitochondrial-enriched proteome. (A) Subcellular localization plot showing proteins from the enriched mitochondrial proteome of Freitas et al. (2022), SVM-predicted mitochondrial proteins identified in this study using a median SVM score cutoff, and proteins shared between the two datasets. A total of 531 proteins were shared between datasets, with 17 proteins uniquely predicted by the SVM classifier and 278 proteins unique to the enriched mitochondrial proteome. (B) Subcellular localization plot showing proteins from the enriched mitochondrial proteome of Freitas et al. (2022), all SVM-predicted mitochondrial proteins identified without application of an SVM score cutoff, and proteins shared between the two datasets. A total of 672 proteins were shared between datasets, with 92 proteins uniquely predicted by the SVM classifier and 137 proteins unique to the enriched mitochondrial proteome. (C) Subcellular localization plot showing the 779 experimentally detected proteins retained in the curated mitochondrial protein compendium. These include 700 proteins supported by both datasets, 66 additional SVM-predicted proteins retained after cluster-based filtering, and 13 proteins retained following evaluation of MitoFates predictions and fractionation profiles. An additional 14 undetected proteins were retained in the final 793-protein compendium.

Proteins not detected in our mass spectrometry dataset were retained only when independent evidence provided additional support for mitochondrial localization. The Freitas et al. mitochondrial compendium combined proteins identified by mass spectrometry with additional proteins incorporated through homology, gene ontology annotation, and previous experimental evidence. Proteins included in their compendium but not detected in our dataset were therefore evaluated for additional evidence supporting mitochondrial localization. Eight proteins with MitoFates scores ≥0.6 were retained, along with two proteins supported by strong homology to established mitochondrial proteins. Four additional proteins encoded by the mitochondrial genome were also retained independently of MitoFates predictions. In total, 14 proteins not detected in our mass spectrometry dataset were incorporated into the curated mitochondrial compendium.

Together, the 700 proteins supported by both datasets, 66 additional SVM-predicted proteins retained after cluster-based filtering, 13 proteins retained following further evaluation (Figure 2C), and 14 proteins retained based on additional mitochondrial evidence produced a curated mitochondrial compendium of 793 proteins (Table S3). These results demonstrate how global subcellular spatial proteomics can both expand and refine existing mitochondrial protein inventories by combining protein fractionation behavior with additional targeting, functional, and genomic evidence.

## Conclusions

This study establishes the first global subcellular localization map of *Dictyostelium discoideum* under vegetative conditions, substantially expanding functional annotation for thousands of previously uncharacterized proteins. Beyond serving as a community resource, this dataset provides a foundation for systematic investigation of protein function, compartmental organization, and cellular architecture.

We are currently preparing manuscripts that further analyze (i) the membrane trafficking system of *Dictyostelium*, including detailed investigation of the contractile vacuole, Jotnarlogs, and patchy proteins, and (ii) the organization of the *Dictyostelium* peroxisome, with particular focus on the subcellular localization of sterol biosynthesis enzymes. Investigators interested in using these data are encouraged to contact us prior to publication to avoid overlap and to facilitate coordinated and collaborative use of this resource.

## Supporting Information

**Figure S1.**
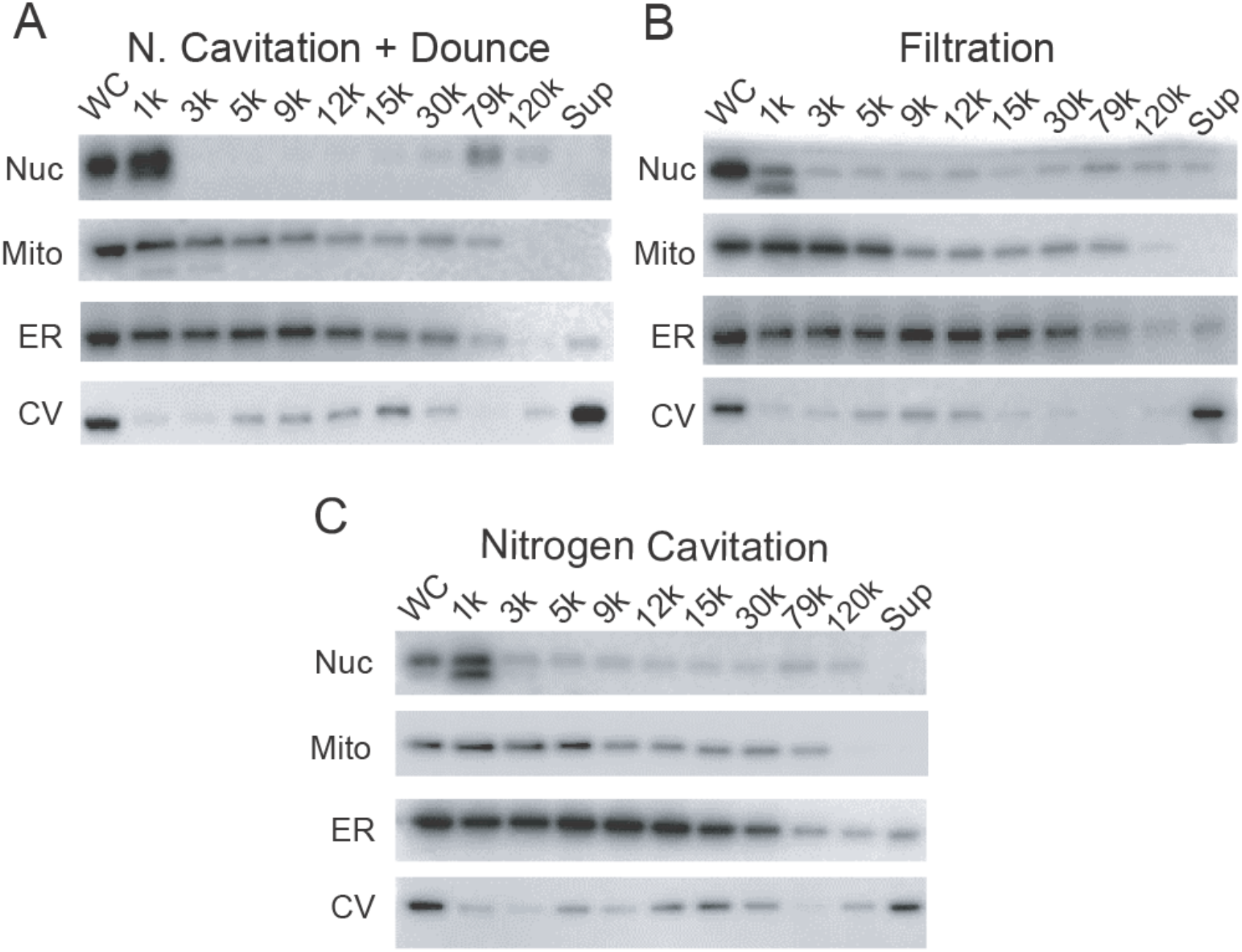
Western blot validation of organelle enrichment across subcellular fractions prior to mass spectrometry. Differential centrifugation fractions, arranged in order of increasing centrifugation speed, were analyzed by Western blot using antibodies against Porin A (mitochondria), V-ATPase subunit A (contractile vacuole), protein disulfide isomerase (PDI; endoplasmic reticulum), and Histone H3 (nucleus). (A) Nitrogen cavitation followed by Dounce homogenization. (B) Filtration. (C) Dounce homogenization only; this method gave poor fractionation and was not carried forward to mass spectrometry. These lysis methods were evaluated as gentle disruption strategies prior to subcellular fractionation and proteomic analysis. The first and last lanes represent whole-cell lysate (WC) and post-ultracentrifugation supernatant (Sup), respectively.

**Figure S2.**
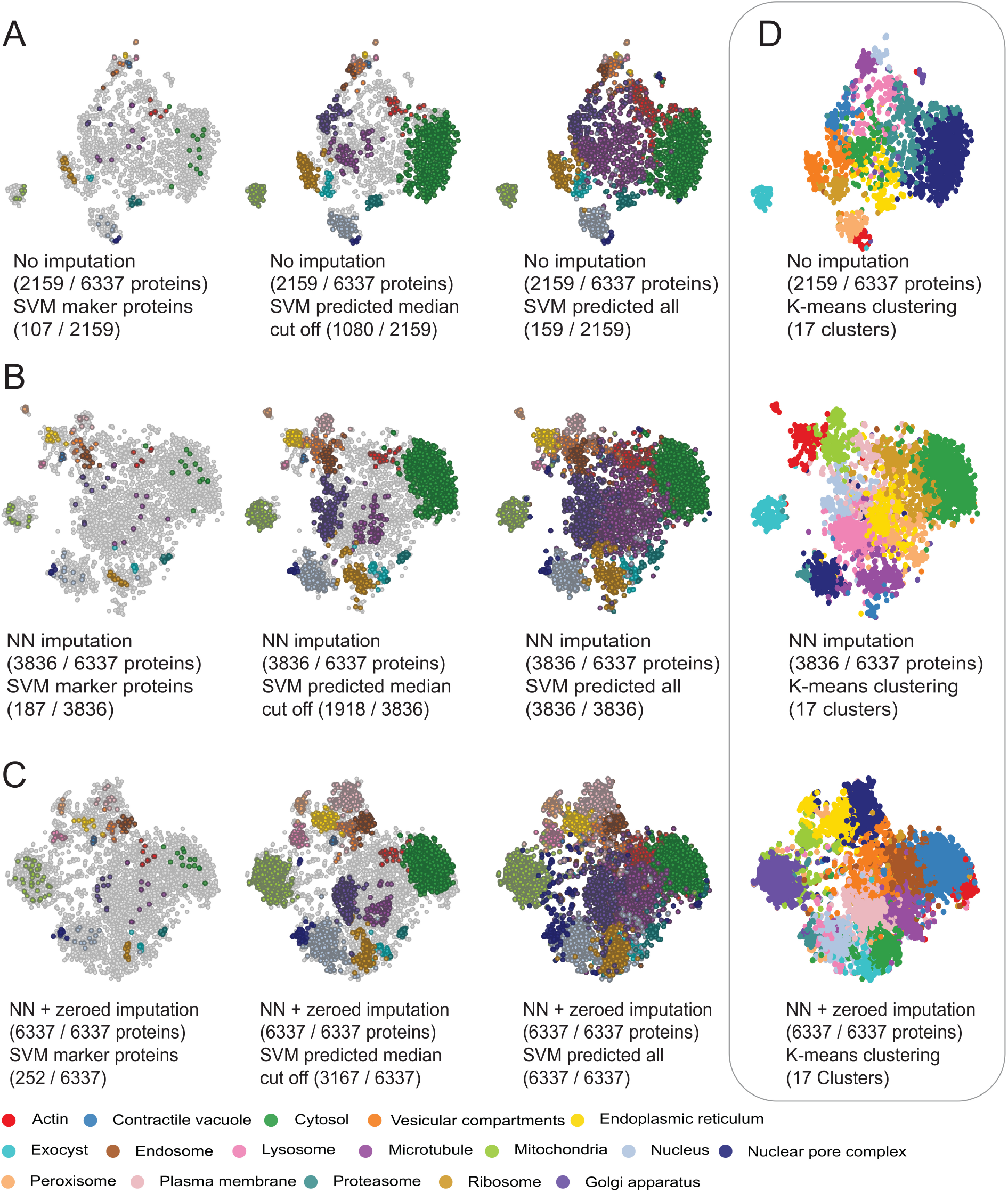
Subcellular atlases of *Dictyostelium discoideum* using different imputation strategies and k-means clustering. Subcellular localization plots generated from the proteomic fractionation dataset using three imputation strategies: (A) no imputation, (B) nearest-neighbor averaging (nbavg) imputation, and (C) nearest-neighbor averaging followed by zero-value imputation. From left to right: marker protein set, SVM median cutoff, and SVM with no cutoff. Colors shown in the accompanying legend correspond to organelle assignments and apply only to the marker protein and SVM-classified datasets. (D) Unsupervised k-means clustering with 17 clusters to match the number of identified subcellular compartments on left. Colors displayed in the k-means clustering panels are used solely to distinguish individual clusters and do not represent organelle identities or correspond to the organelle color scheme used in the supervised classifications.

**Table S1:** Curated marker set, comprising proteins that are either experimentally localized in Dictyostelium discoideum or have strong homology to validated markers in other eukaryotes, with the 252 proteins used to train the SVM classifier indicated.

**Table S2:** SVM prediction values for both no cut-off and median cut-off.

**Table S3:** Curated mitochondrial protein compendium

## Methods

### Culturing and Cell Harvesting

*Dictyostelium discoideum* strain DBS0237699 (AX2) was obtained from the Dicty Stock Center at Northwestern University and cultured using axenic techniques described by (Fey et al., 2006; Watts and Ashworth, 1970). Cells were inoculated at approximately 5 × 10^4^ cells mL^−1^ into 1,000 mL Erlenmeyer flasks containing 200 mL of HL5 medium (Bacto Peptone, yeast extract, maltose monohydrate, streptomycin) and maintained at room temperature with shaking at 180 r.p.m. until reaching a density of 1 × 10^6^ cells mL^−1^. Cell density was monitored using a hemocytometer. Cultures were harvested by centrifugation at 1,100 × g for 2 min at 4°C and washed twice with detergent-free (DF) buffer (0.25 M sucrose, 10 mM HEPES pH 7.4, 2 mM EDTA, 2 mM magnesium acetate). The resulting cell pellet was resuspended in 3 mL of ice-cold DF buffer, and Halt™ Protease and Phosphatase Inhibitor Cocktail (Thermo Scientific) was added immediately prior to lysis.

### Cell Lysis and Fractionation

Three different lysis conditions were employed to gently disrupt *Dictyostelium* cells while preserving subcellular compartments.

#### Nitrogen Cavitation

The nitrogen cavitation chamber was prepared by cleaning with 70% ethanol and equilibrating on ice with DF buffer. Washed cells were added into the pre-chilled chamber and pressurized with ultra-pure nitrogen gas at 1,000 psi for 10 min on ice. Lysate was collected dropwise by slowly releasing the outer valve into a pre-chilled tube.

#### Filtration

Filtration was performed following the method of (Prem Das and Henderson, 1983). Washed cells were loaded into a 10 mL syringe fitted with a filtration chamber containing two stacked Avantor™ Nucleopore filters (5.0 µm pore diameter). Cells were lysed by forced passage through the filters and the filtrate was collected into a pre-chilled tube.

#### Nitrogen Cavitation Followed by Dounce Homogenization

Cells were first lysed by nitrogen cavitation as described above. The resulting lysate was then transferred to a pre-chilled Teflon B-grade Dounce homogenizer and further disrupted by 12 gentle strokes on ice.

Following lysis, 1 mL of cell lysate from each lysis method was divided into two tubes. Subcellular fractionation was performed according to the differential centrifugation protocol described previously (Geladaki et al., 2019). One set of fractions was reserved for Western blot analysis, while the other set was stored at –80°C for downstream mass spectrometry.

### Western Blofling

Western blots were performed using four different antibodies. These assays were used to verify that the gentle lysis conditions preserved subcellular structures. They also confirmed that subcellular proteins were successfully separated into distinct fractions during fractionation.

Pellets obtained from differential centrifugation were resuspended in 2X Laemmli buffer (4% SDS, 20% glycerol, and 120 mM Tris-HCl pH 6.8) and heated at 95 °C until pellets dissolved. Resuspended proteins were quantified using Pierce™ BCA Protein Assay Kit (Thermo Scientific) according to the manufacturer’s instructions. 5 µg of proteins supplemented with bromophenol blue were loaded onto Invitrogen™ Bolt™ Bis-Tris Plus Mini Protein Gels (4–12%, 1.0 mm, WedgeWell™ format) alongside the Amersham™ ECL™ Rainbow™ Marker (Full Range). Electrophoresis was performed at 125 V for 1h. Proteins were transferred to PVDF membranes using the iBlot™ 2 Transfer Stacks (Invitrogen™) and the iBlot™ 2 Western Blot Transfer Device following the default program optimized for Bolt™ gels (25V for 6 minutes). Membranes were activated with methanol and briefly rinsed with deionized water and blocked for 1 h at room temperature in Kroger™ 5% dry milk prepared in 1× Tris-buffered saline (TBS) with gentle shaking. Primary antibody incubation was performed using four rabbit-derived antibodies: anti-Mitochondrial outer membrane porin (ABCD_AK421; monoclonal), anti-V-ATPase subunit A (ABCD_AJ520; monoclonal), anti-Protein disulfide-isomerase 1 (ABCD_AN703; monoclonal), and anti-Histone H3 (Abcam, AB1791; polyclonal). Primary antibodies were diluted in blocking buffer (1:1,000 for ABCD antibodies and Histone), incubated for 1 h at room temperature with gentle rocking, and then overnight at 4°C. Following three 10-min washes in TBS, membranes were incubated for 1 h at room temperature with Goat anti-Rabbit IgG (H&L) HRP-conjugated secondary antibody (ImmunoReagents Inc.; 1:1,000 in blocking buffer), washed three times for 10 min each in TBS-T (TBS + 0.05% Tween-20), and briefly rinsed in water. Protein bands were detected using the Pierce™ ECL Western Blotting Substrate (Thermo Scientific) and visualized using an Azure 600 Biosystems chemiluminescence imager.

### Proteomics of Dictyostelium Organellar Fractions by LC-MS/MS

Pelleted organellar fractions were resuspended in 500 µL lysis buffer (8M urea, 75mM NaCl, 1mM EDTA, 1x HALT, 50mM Tris-HCl at pH 8.0), and sonicated for 1 minute with four 15-second cycles at 50% amplitude. The samples were then centrifuged at 20,000 rcf at 4°C for 10 min. The protein supernatant was quantified by BCA (Thermo). From each sample, 70 µg of protein was reduced with 5 mM dithiothreitol (DTT), alkylated with 20 mM iodoacetamide (IAA) and digested overnight with trypsin at an 1:25 enzyme:substrate ratio (Trypsin Gold, MS grade, Promega), at 37°C. After overnight digestion, peptides were desalted using 1cc C18 SPE columns (Waters Seppak), vacuum-concentrated to dryness and stored at −80°C until analysis. Desalted peptides were then reconstituted in 100 µL of 2% acetonitrile, 0.1% formic acid in water, and quantified by BCA protein assay. To generate a spectral library, equal amounts of peptides from each sample were pooled and fractionated to 24 fractions by HPLC high-pH RP fractionation. Peptide fractions were vacuum-concentrated to dryness. All samples and library fractions were reconstituted in 2% acetonitrile, 0.1% formic acid in water containing iRT peptides (Biognosys). LC-MS/MS data were acquired using a Vanquish Neo ultra high-performance liquid chromatography (UHPLC) system (Thermo Fisher Scientific) coupled to an Orbitrap Ascend Tribrid mass spectrometer (Thermo Fisher Scientific) equipped with a FAIMS Pro device. Peptides (5 µL injections) were directly loaded on a C18 analytical column (EasySpray ES802A, 75 µm inner diameter × 25 cm, 2-µm particle size, 100 Å pore size) kept at 45°C, and separated at a flow rate of 300 nL/min. Peptides were eluted from the column using a 120 min gradient formed by Solvent A (LC-MS grade Water, 0.1% formic acid) and Solvent B (LC-MS grade acetonitrile, 0.1% formic acid). Library fractions were acquired in DDA mode using the following settings: MS1 in Orbitrap at 120K resolution, MS2 in Orbitrap at 30K resolution, HCD NCE of 25%. All samples were acquired in DIA mode using the following settings: DIA scan over a mass range of 380-980 m/z, isolation window of 16 m/z with 1 m/z overlap, detection in the Orbitrap at 15K resolution, HCD NCE of 25% and FAIMS CV at −45. A sample-specific spectral library was generated by Pulsar search engine in Spectronaut (v19.0.240606.62635) with the following parameters: mapped against the *Dictyostelium discoideum* (dicty_2.7) proteome, peptide modifications set for carbamidomethyl and oxidation, allowed for two missed cleavages and proteins and peptides filtered for a false discovery rate less than 1%.

DIA-MS data were searched against the sample-specific library in Spectronaut (v19.0.240606.62635) using default settings, except that cross normalization was turned “OFF”. Results were filtered to 1% FDR for high confidence protein and peptide identification.

Protein and peptide abundances generated from Spectronaut were normalized using variance stabilizing normalization in R (v4.3.0).

### Raw data processing

Individual protein abundance profiles from each of the three lysis conditions were merged to generate a comprehensive protein abundance dataset spanning all fractions. Proteins that were completely absent in any condition were excluded from further analysis. The remaining proteins and their corresponding abundance values were imported into R and converted into an MSnSet object using the Bioconductor packages MSnbase (v2.20.1; Gatto and Lilley, 2012) and pRoloc (v1.34.0; Gatto et al., 2014). The resulting dataset, which already included imputed values, was used for downstream analysis. t-Distributed Stochastic Neighbor Embedding (t-SNE) was then applied for dimensional reduction and visualization of the protein distribution across subcellular fractions (van der Maaten and Hinton, 2008).

### Supervised Classification

SVM classification was performed on post-imputation marker protein abundance profiles using the Bioconductor pRoloc package (v1.34.0) in R. A curated marker set of 415 proteins across 17 subcellular compartments was assembled (Table S1), from which 252 proteins were selected and used to train the classifier. The optimal SVM parameter**s** determined from the marker set were then applied to all proteins in the dataset, generating SVM scores ranging from 0 to 1, with 1 corresponding to marker protein confidence. Proteins without labels (non-marker proteins) were classified using these parameters, with weights applied according to the marker classes. Proteins with SVM scores below the global median were reset to “unknown,” while proteins above the median were considered predicted to their corresponding compartments.

### Targeting sequence identification

SignalP 6.0 was first applied to identify signal peptides indicative of Sec/SPI-mediated targeting (Teufel et al., 2022). DeepTMHMM v1.0.24 (Hallgren et al., 2022) was then used to predict transmembrane topology, and MitoFates v1.2 (Fukasawa et al., 2015) was employed to detect mitochondrial-targeting signals, with proteins scoring above 0.9 classified as mitochondrial-targeted. Nuclear localization signals were predicted using NLStradamus r.9 (Nguyen Ba et al., 2009), and proteins with prediction scores greater than 0.95 were classified as containing a nuclear localization signal. Predicted targeting features were mapped onto the subcellular localization plot to examine their distribution across cellular compartments.

### Establishment of a Curated Mitochondrial Protein Compendium

The curated mitochondrial protein compendium was generated by integrating our SVM-based mitochondrial predictions with the Freitas et al. (2022) mitochondrial compendium. Both median-cutoff and no-cutoff SVM predictions were compared with the Freitas dataset. Using the no-cutoff predictions, 672 proteins were shared by both datasets. An additional 28 Freitas proteins that were not predicted as mitochondrial by the SVM but localized adjacent to the mitochondrial cluster were retained after inspection of their fractionation profiles, resulting in 700 proteins supported by both datasets.

Our SVM classifier also predicted 92 mitochondrial proteins absent from the Freitas et al. compendium. These proteins were evaluated based on their positions relative to the mitochondrial cluster. Proteins outside the cluster were excluded, resulting in the retention of 66 additional proteins. Of the 109 proteins from the Freitas et al. compendium that were detected in our mass spectrometry data but assigned outside the mitochondrial cluster, 13 were retained after further evaluation of their MitoFates predictions and fractionation profiles. Three had MitoFates scores from 0.3 to less than 0.6, and 10 had scores ≥0.6. These proteins were retained when their profiles resembled those of mitochondrial proteins in one or more fractionation experiments.

Proteins present in the Freitas et al. compendium but not detected in our dataset were evaluated separately. Eight proteins with MitoFates scores ≥0.6, two proteins with strong homology to established mitochondrial proteins, and four mitochondrial genome-encoded proteins were retained. Together, these criteria produced a curated mitochondrial protein compendium of 793 proteins (Table S3).

## Data Availability

The mass spectrometry proteomics data have been deposited to the ProteomeXchange Consortium via the PRIDE partner repository with the dataset identifier PXD084378.

## Acknowledgments

J.G.W. was supported by a grant from the National Science Foundation (DBI-2119963) and a grant from the Gordon and Betty Moore Foundation (GBMF10600). This research includes work conducted in the Integrated Mass Spectrometry Shared Resource supported by the National Cancer Institute of the NIH under grant (P30CA033572).

